# LRRK2 G2019S disrupts GABAergic signaling and shifts excitatory/inhibitory balance in the striatum

**DOI:** 10.1101/2025.10.23.684189

**Authors:** Angela Di Iacovo, Giuseppe Aceto, Ludovica Iovino, Roberta De Rosa, Raffaella Cinquetti, Charlotte Kilstrup-Nielsen, Tiziana Alberio, Marcello D’Ascenzo, Laura Civiero, Elena Bossi, Cristina Roseti

## Abstract

The excitatory/inhibitory (E/I) balance within neural circuits is essential for proper brain function, and its disruption is a hallmark of several neurodegenerative diseases. In Parkinson’s disease (PD), widespread alterations in the basal ganglia circuitry lead to an E/I imbalance in the striatum, contributing to excitotoxicity.

Leucine-rich repeat kinase 2 (LRRK2) has recently emerged as a key contributor to both familial and sporadic forms of PD, with the pathogenic Gly2019Ser (G2019S) mutation representing one of the most frequently observed variants. This mutation is known to exacerbate excitotoxicity by impairing glutamate reuptake mechanisms, particularly through dysregulation of EAAT2 activity and its membrane localization. In contrast, the role of LRRK2 in GABAergic transmission remains poorly understood. Here, we reveal a clear modulation of inhibitory signaling by LRRK2 through a comprehensive approach combining mouse striatal slices and *Xenopus laevis* oocytes. Our results demonstrate, for the first time, that LRRK2 G2019S induces a significant reduction in GABA-evoked current amplitudes. Moreover, we identified an altered distribution of receptor isoforms in pathological tissue, affecting both tonic and phasic GABA currents. Specifically, synaptic GABA_A_ receptors containing the γ_2_ subunit were functionally modulated by LRRK2 G2019S. The reduced availability of gephyrin in the presence of the G2019S variant may impair the gephyrin–GABA_A_ receptor complex, leading to decreased receptor surface expression and further shifting the glutamate/GABA current ratio toward excitatory dominance. This is supported by the increased activity of AMPA and NMDA receptors observed in the pathological striatum.

Overall, our findings highlight a previously underappreciated role of LRRK2 G2019S in impairing GABAergic transmission and disrupting the E/I balance. These insights point to novel circuit-level mechanisms underlying LRRK2-linked PD and suggest new avenues for the development of disease-modifying therapies targeting inhibitory dysfunction.

**GRAPHICAL ABSTRACT:** 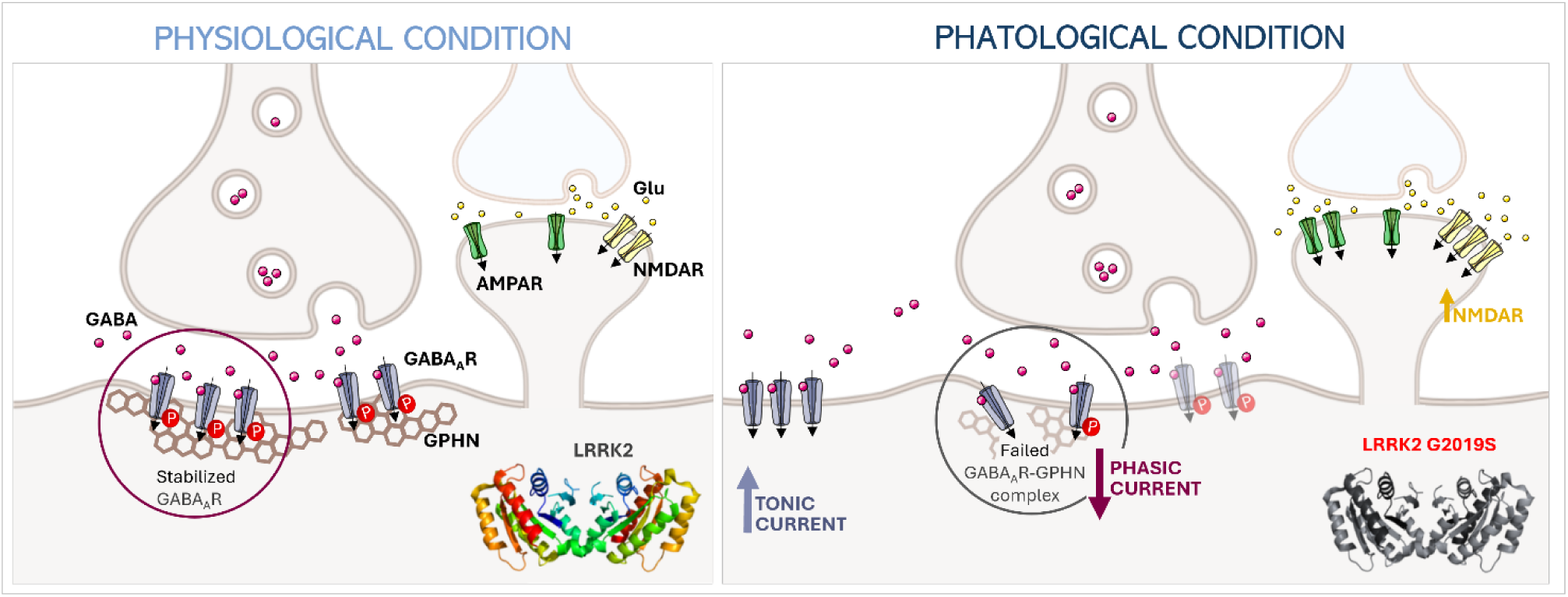

The LRRK2 G2019S mutation contributes to excitatory/inhibitory imbalance by reducing GABA-evoked currents. Specifically, it is associated with diminished phasic GABAergic transmission and enhanced tonic inhibition, suggesting an altered subcellular distribution of GABA_A_ receptor subtypes.

## Introduction

Mutations in the LRRK2 gene are found at a high frequency in Parkinson’s disease (PD) patients, with the G2019S substitution being the most common. It leads to increased LRRK2 kinase activity.^1^ Interestingly, patients with LRRK2-associated PD have shown clinical features almost indistinguishable from the sporadic form of the disease, suggesting that there could be a central role of LRRK2 in the pathogenesis of both forms of PD.^2^

*LRRK2* encodes for a large multi-protein enzyme consisting of 2527 amino acids that is largely distributed in the brain, particularly in neurons, microglia, and astrocytes.^3^ LRRK2 is physically and functionally associated with many components in the neuronal cells, taking part in many physiological cellular processes.^4,5^ Over the years, it has been demonstrated that LRRK2 G2019S causes deficits in vesicular trafficking and endo-lysosomal system.^6^ In particular, the pathological LRRK2 G2019S variant disrupts glutamate homeostasis by impairing the membrane localization of EAAT2, thereby reducing astrocytic glutamate reuptake capacity and contributing to synaptic glutamate dyshomeostasis.^7^Moreover, the cortical neurons from LRRK2 G2019S knock-in (KI) mice showed increased miniature and spontaneous excitatory postsynaptic current (mEPSC and sEPSC) frequency,^8,9^ indicating a correlation between aberrant cortico-striatal glutamatergic neurotransmission and the mutated form of LRRK2 in PD.

The majority of striatal neurons are GABAergic medium spiny neurons (MSNs), which express either dopamine receptor type 1 (D_1_-R) or type 2 (D_2_-R). These neurons receive dopaminergic input from the *substantia nigra pars compacta* and are key components of the direct and indirect pathways that regulate motor control. Moreover, the activity and synchronization of MSNs are locally modulated by different subtypes of GABAergic and cholinergic interneurons. LRRK2 is expressed in MSNs, highlighting its potential involvement in modulating GABAergic transmission and, consequently striatal output.^5^

Thus, since GABA is the main inhibitory neurotransmitter within the striatum, GABA_A_ receptor - mediated currents are determinant for striatal activity, maintaining the optimal levels of neuronal excitability.^10^ GABA_A_ receptors are ligand-gated chloride channels composed of heteromeric pentamers formed by various combinations of α (α_1_-_6_), β (β_1_-_3_), γ (γ_1_-_3_), δ, ε, π, θ, and ρ subunits. Based on their subunit composition, they are located in the synapses or extrasynaptically. Generally, GABA_A_ receptors containing the γ-subunit are located at synapses, where they respond to high transient concentrations of GABA and mediate phasic inhibition. On the other hand, δ-subunit containing receptors are found at extrasynaptic and perisynaptic locations and cause tonic inhibition; indeed, extra-synaptic GABA_A_ receptors take part in long-run modulation of neuronal activity after GABA spillover or non-synaptic GABA release.^11^ Because of the importance of GABAergic signalling, several mechanisms finely regulate dynamic adaptation of cell surface expression, endocytosis and recycling of GABA_A_ receptors. Direct and indirect interactions with the postsynaptic scaffold act in regulating their membrane localization to maintain the synaptic activity.^12^

While substantial evidence has established a role for LRRK2 in modulating excitatory signaling in PD, its impact on GABAergic transmission remains largely unexplored. Given that GABAergic signaling is critical for maintaining the balance of glutamatergic neuronal excitability, the aim of our study was to investigate the role of LRRK2 in striatal GABAergic transmission. In particular, using a combination of electrophysiological and molecular approaches, we investigated the effects of the pathogenic LRRK2 G2019S variant on the functional properties and membrane expression of GABA_A_ receptors.

## Materials and methods

### Samples preparation: striatal membranes, cRNAs and DNAs

C57Bl/6J Lrrk2 wild-type (WT) and Lrrk2 G2019S knock-in mice striata were used (4-months old). All procedures performed with mice were approved by the Ethical Committee of the University of Padova and the Italian Ministry of Health (license 200/2019). Using a Teflon glass homogenizer, about 0.5 g of previously frozen tissue was homogenized in 2 ml of glycine buffer (200 mM glycine/150 mM NaCl/50 mM EGTA/50 mM EDTA/300 mM sucrose), plus 20 μl of protease inhibitors (Sigma P2714), pH 9 with NaOH. The filtrate was centrifuged for 15 min at 9,500 × g in a Beckmann centrifuge (C1015 rotor). The supernatant was then centrifuged for 2 h at 100,000 × g in a SW40 rotor at 4°C in a Beckmann centrifuge. The pellet was washed, resuspended in 5 mM glycine, aliquoted and kept at −80°C for later use.^13^ The recombinant plasmids coding for LRRK2 WT and G2019S, and GABA_A_ receptor subunits were introduced into JM109 strain of *E. coli* after heat-shock procedure. Capped cRNAs were synthesized by in vitro transcription using T7 RNA polymerase from cDNAs linearized with restriction enzymes and then purified. Data about the plasmid vectors used for heterologous expression in X. laevis oocytes are detailed in Table 1.

**Table 1.** List of genes and vectors characteristics used for heterologous expression in X. laevis oocytes; (*LRRK2* WT and G2019S cDNAs were linearized using restriction enzyme SmaI).

| Protein | Gene | Organism | Vector |
| --- | --- | --- | --- |
| LRRK2 WT | *LRRK2* | Homo sapiens | pDESTN-SF-TAP |
| LRRK2 G2019S | *LRRK2* | Homo sapiens | pDESTN-SF-TAP |
| α1 subunit | *Gabra1* | Rat | pGW1 |
| β2 subunit | *Gabrb2* | Rat | pGW1 |
| γ2 subunit | *Gabrg2* | Rat | pGW1 |

### Injection procedures in *Xenopus laevis* oocytes

The experiment with *Xenopus laevis* oocytes was conducted using an experimental protocol approved locally by the Ethical Committee of the Organismo Preposto al Benessere degli Animali (OPBA) of the University of Insubria (OPBA permit no. 06_20) and by the Italian Ministry of Health (permit no. 440/2021-PR). Oocytes were obtained from adult *Xenopus laevis* females. Frogs were anaesthetised by immersion in 0.1% (w/v) MS222 (tricaine methanesulfonate; Sigma) solution in tap water adjusted at final pH 7.5 with bicarbonate. Abdomens were sterilized with antiseptic agent (Povidone-iodine 0.8%) and ovary portions were removed by laparotomy. The collected oocytes were treated with 0.5 mg/ml collagenase (Sigma Collagenase from Clostridium histolyticum) in calcium-free ND96 (96 mM NaCl, 2 mM KCl, 1 mM MgCl_2_, 5 mM 4-(2-hydroxyethyl)-1 piperazineethanesulfonic acid (HEPES); pH 7.6) for at least 1 h at 18°C.

Healthy and fully grown oocytes, appearing to be at stage V and VI, were selected and manually separated in NDE solution (ND96 plus 2.5 mM pyruvate, 0.05 mg/mL gentamicin sulphate and 1.8 mM CaCl_2_).^14^ The following day, the oocytes were injected with the experimental samples using a manual microinjection system (Drummond Scientific Company, Broomall, PA).

A volume of 80 nl mouse striatal membranes was injected per oocyte. The oocytes were ready for current recordings 24 h after injection. The purified cRNAs (LRRK2 WT and G2019S) were injected at a concentration of 25 ng/µl per oocyte and tested after 72 h. cDNAs were injected into the nucleus at a concentration of 160 ng/µl, 48 h before testing. When cRNA and membrane fragments were co-expressed in oocytes, the membrane samples were injected 48 h after the cRNA injection. Prior to electrophysiological recordings, the oocytes were incubated at 18 °C in NDE solution.

### Electrophysiological Recordings from Oocytes

Electrophysiological studies were performed using the Two-Electrode Voltage Clamp (TEVC) technique (Oocyte Clamp OC-725; Warner Instruments, Hamden, CT, United States). Borosilicate microelectrodes, with a tip resistance of 0.5–4 MΩ, were filled with 3 M KCl. Bath electrodes were connected to the experimental oocyte chamber via agar bridges (3% agar in 3 M KCl). The external control solution ND98 had the following composition: 98 mM NaCl, 1 mM MgCl_2_, 1.8 mM CaCl_2_, 5 mM HEPES, adjusted to pH 7.6 with NaOH. Signals were filtered at 0.1 kHz and sampled at 200 Hz or 0.5 kHz and at 1 kHz.

The experiments were performed using each substrate at saturating concentration to record the current at -60 mV. The AMPA receptor current was elicited by applying the allosteric modulator, cyclothiazide (CTZ 10 µM), for 20 s as pretreatment before the perfusion of the substrate α-Ammino-3-idrossi-5-Metil-4-isossazol-Propionic Acid (AMPA 20 µM). The NMDA receptor-current was recorded using glutamate (Glu 200 µM) with Glycine (10 µM) and Serine (10 µM) co-perfusion at Vh -50 mV. In this last case, we used the extracellular recording solution Mg^2+^-free and Ca^2+^-free (100 mM NaCl, 0.3 mM BaCl_2_, 5 mM HEPES, pH 7.2 adjusted with KOH) in order to avoid contamination of the NMDA receptor current by the endogenous Ca^2+^-activated Cl^−^ conductance ^15^. To discriminate between NMDA and AMPA evoked currents, the oocytes were treated with selective and potent antagonist of NMDA receptor, Kynurenic acid (KYNA, 500 µM), and inhibitor of AMPA receptor and Kainate receptor, 2,3-Dioxo-6-nitro-1,2,3,4-tetrahydrobenzo[f]quinoxaline-7-sulfonamide (NBQX, 10 µM). To evaluate GABA reversal potential (E_GABA_), the ramp protocol from -120 mV to +40 mV lasting for 920 ms was applied. The resulting current-voltage (I-V) relationship was determined by subtracting the current recorded in ND98 buffer from the evoked substrate current.

GABA_A_ receptor dose-current response relation was obtained using increasing concentrations of GABA (1 µM, 10 µM, 30 µM, 100 µM, 300 µM, 1000 µM, 3000 µM). GABA currents were normalized to the maximal current (I_max_) for each tested cell. To calculate the kinetic parameters, the fitting of GABA_A_Rs dose-response was fitted with Hill equation. LRRK2 kinase inhibitor MLi-2 was used at 200 nM concentration for 90 min application; (Tocris; Cat. #5756).^716^

### Electrophysiological Recordings from mice brain slices

To prepare brai2n slices, we followed the protocol described by Aceto et al. 2022 (Journal of Physiology), with minor modifications. Male C57BL/6J mice (2 months old) were used from colonies bred at the animal facilities of Università Cattolica del Sacro Cuore (authorization no. 623/2022-PR). Animals were sacrificed by cervical dislocation. Brains were rapidly removed and placed in ice-cold cutting solution containing (in mM): TRIS-HCl 72, TRIZMA base 18, NaH_2_PO_4_ 1.2, NaHCO_3_ 30, KCl 2.5, glucose 25, HEPES 20, MgSO_4_ 10, Na-pyruvate 3, ascorbic acid 5, CaCl_2_ 0.5, sucrose 20. Slices (300 μm thick) were cut on a vibratome (VT1200S; Leica Microsystems, Germany) and immediately transferred to an incubation chamber held at 32°C and filled with a recovery solution containing (in mM): TRIS-HCl 72, TRIZMA base 18, NaH_2_PO_4_ 1.2, NaHCO_3_ 25, KCl 2.5, glucose 25, HEPES 20, MgSO_4_ 10, Na-pyruvate 3, ascorbic acid 5, CaCl_2_ 0.5, sucrose 20. After 30 min, slices were transferred to a second incubation chamber held at 32°C and filled with artificial cerebrospinal fluid (aCSF) containing (in mM): NaCl 124, KCl 3.2, NaH_2_PO_4_ 1.2, MgCl_2_ 1, CaCl_2_ 2, NaHCO_3_ 26, and glucose 10, pH 7.4. During incubations, the chambers were continuously bubbled with 95% O_2_/5% CO_2_. Slices were equilibrated at room temperature for at least 45 min. Slices were then transferred to a submerged recording chamber constantly perfused with heated aCSF (32°C) and bubbled with 95% O_2_/5% CO_2_. MSNs within the dorsal striatum subregion were identified with a 40X water-immersion objective on an upright microscope equipped with differential interface contrast optics under infrared illumination (BX5IWI, Olympus, Center Valley, PA) and video observation. Electrodes were made from borosilicate glass micropipettes (Warner Instruments, Hamden, CT) prepared with a P-97 Flaming-Brown micropipette puller (Sutter Instruments, Novato, CA). Patch pipettes had a resistance of 4-6 MΩ when filled with an internal solution containing (in mM): CsCl 135, HEPES 10, EGTA 1.1, CaCl_2_ 0.1; Mg-ATP 2.5, Na-GTP 0.25, phosphocreatine 5, pH 7.2. After establishing a gigaseal, the patch was broken by applying negative pressure to achieve a whole-cell configuration. A series resistance lower than 15 MΩ was considered acceptable and monitored constantly throughout the entire recording. Neurons were held at -70 mV. Tetrodotoxin (TTX, 0.5 μM, Tocris), D-(-)-2-Amino-5-phosphonopentanoic acid (D-AP5, 50 μM, Tocris) and NBQX (10 μM, Tocris) were applied to the bath to block action potential-mediated neurotransmitter release, NMDA and AMPA receptors, respectively. MLi-2 (200 nM) was intracellularly perfused through the patch-pipette. Recordings were performed using a Multiclamp 700B/Digidata 1550A system (Molecular Devices, Sunnyvale, CA) and digitized at a 10,000 Hz sampling frequency. Electrophysiological recordings were analyzed using the Clampfit 10.9 software (Molecular Devices). A template was constructed using the “Event detection/create template” function, as previously described,^17,18^ then, miniature inhibitory postsynaptic currents (mIPSCs) were detected using the “Event detection/template search” function. All the waveforms detected during a single recording using template analysis were averaged and amplitude, rise time and decay time calculated. The amplitudes, frequencies, and the kinetics of mIPSCs were measured after 10 min of intracellular dialysis.

### Immunoblotting

Striatal samples derived from WT and LRRK2 G2019S knock-in mice were lysed in RIPA buffer containing 1% protease inhibitor cocktail (Sigma-Aldrich). The protein concentration of total striatal lysates and striatal membrane fragments derived from LRRK2 WT and LRRK2 G2019S KI mice was measured using the Pierce® BCA Protein Assay Kit following the manufacturer’s instructions (Thermo Scientific). Samples were transferred to nitrocellulose membranes and blocked in 5% non-fat milk in TBS (20 mM Tris-HCl, pH 7.5, 150 mM NaCl) with 0.2% Tween-20 (T-TBS). Blots were incubated with primary antibodies overnight at 4 °C, washed in T-TBS, and incubated with appropriate secondary antibody for 1 h at room temperature. After extensive washes blots were developed with Protein Detection System-ECL (Genespin) coupled to G:BOX Chemi Imaging System (Syngene). Antibodies used: rabbit anti-GABA_A_R γ_2_ (SYSY, 224003), rabbit anti-GAPDH (Sigma, G9545), mouse anti-gephyrin (Santa Cruz, sc-25311). Secondary Antibodies: HRP-conjugated goat anti-mouse, goat anti-rabbit and donkey anti-goat secondary antibodies for western blottings were purchased from Jackson Immunoresearch. GAPDH signal and Ponceau loading were used as normalizers for our protein of interest in total lysates and membranes samples, respectively.

### Single oocyte chemiluminescence (SOC)

The oocytes were fixed in ice-cold 4% PFA in PBS pH 7.5, for 15 min at 4 °C. After fixation the oocytes were subjected to a 1 h incubation in a blocking solution (1% BSA in ND96) and then incubated for 2 h with primary antibody mouse anti-gephyrin (1:250 in blocking solution) at 4 °C, followed by a 5 min wash repeated six times. The oocytes were then transferred RT and maintained for 1 h in peroxidase-conjugated anti-mouse (1:1000 in blocking solution), with rinsing for 3 min repeated six times in 1% BSA in ND96 and 3 min repeated six times in ND96 alone. Finally, each oocyte was transferred to 96-well plate containing 50 μl of SuperSignal West Femto (Pierce, Thermo Fisher Scientific). The chemiluminescent signal was quantified using a Tecan Infinity 200 microplate reader (Tecan). The plate was read within 5 min after the transfer of the first oocyte.^16^

### Protein Networks

Cytoscape 3.10.1 was used to generate a PPI network connecting γ_2_ subunit of GABA_A_ receptor (GABRG2), Gephyrin (GPHN) and LRRK2 to investigate their physical interactions and common interactors.^19^ The public database of the International Molecular Exchange Consortium (IMEx) was queried through Cytoscape using the PSICQUIC standard (the Proteomics Standard Initiative Common QUery InterfaCe).^20^ The network was filtered for interactor type protein, and for taxonomy ID 9606 (Homo sapiens), 10116 (Ratus norvegicus), and 10090 (Mus musculus). Duplicated edges and self-loops were removed. Only proteins connected in the main network were retained. Common interactors to at least two proteins among GABRG2, GPHN and LRRK2 have been selected with their first interactors and the related sub-network extracted. A new network was built starting from this new input list (26 nodes) to retrieve information about reciprocal interactions, second interactors have been removed. The significance of the PPI network was assessed using permutation testing. The network was randomized multiple times (1000 iterations) with the same number of nodes, but edges assigned randomly. The observed node/edge number ratio was compared to the randomized distribution. The p-value was computed as 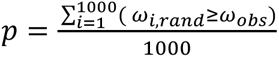.

### Data analysis and statistics

All electrophysiological data analysis was collected using Clampfit 10.7 software (Molecular Devices, Sunnyvale, CA, USA) and analyzed with OriginPro 8.0 (OriginLab Corp., Northampton, MA, USA). GraphPad Prism 8.0.2 (GraphPad Software, Boston, MA, USA) was used for statistical analysis and figures preparation. Immunoblotting quantification analysis was performed by ImageJ. We assessed the normality of the data using the Shapiro–Wilk test to determine whether to apply parametric or non-parametric (Wilcoxon signed-rank test, Mann–Whitney U test) tests before proceeding with the analysis. Statistical analysis of the data was performed with the unpaired *t*-test or Mann–Whitney test within two different experimental groups. A paired *t*-test (or Wilcoxon signed-rank test) was applied for the comparison of two conditions of the same group. Levels of significance were defined as *, p<0.05; **, p <0.01; ***, p <0.001; ****, p <0.0001. The number of oocytes/slices (*n*) obtained from a frog/mouse (*N*) used in each experiment is indicated as “*n/N*”.

## Results

### LRRK2 inhibitor decreases GABA_A_-mediated currents in striatal MSNs

We first assessed the potential involvement of LRRK2 WT in regulating GABAergic signalling in striatal slices from WT mice (Fig. 1). To pharmacologically modulate LRRK2, we used the highly selective LRRK2 inhibitor MLi-2 (200 nM) that was injected into the recorded MSNs via a patch pipette. Under these experimental conditions the peak amplitudes of mIPSCs were significantly decreased compared to what was observed following intracellular perfusion with vehicle (DMSO; 1/1000; I_vehicle_: 29.5 ± 1.9 pA *vs* I_MLi-2_: 19.6 ± 2.7 pA; Unpaired Student’s t-test, *p*=0.003; Fig.1A-1C). In contrast, there was no significant difference in mIPSC frequency (Vehicle: 1.4 ± 0.1 Hz *vs* MLi-2: 1.2 ± 0.2 Hz; Unpaired-t test; *p*>0.05; Fig. 1D) and mIPSC kinetics (Rise time: vehicle = 1.7 ± 0.1 pA *vs* MLi-2 =1.6 ± 0.2 pA; Unpaired-t test; *p*>0.05; decay time: vehicle=14.4 ± 0.5 ms *vs* MLi-2=13.1 ± 0.7 ms; Unpaired-t test; p>0.05; Fig. 1E-1F). These results suggest that the activity of LRRK2 influences inhibitory synaptic transmission of dorsal striatum MSNs.

**Figure 1.**
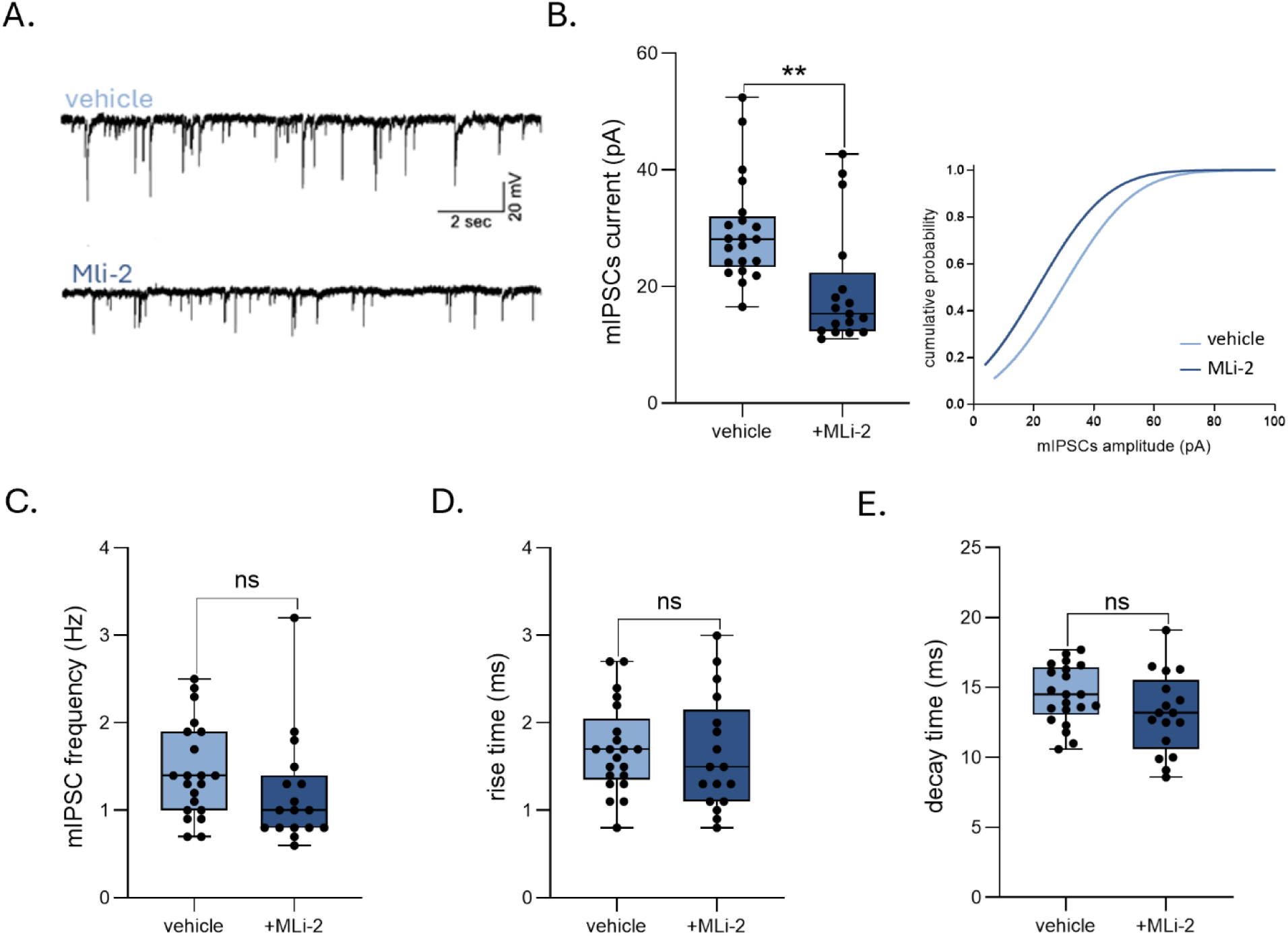
Postsynaptic inhibition of LRRK2 decreases GABA_A_ currents in MSNs. (**A)** Representative traces of mIPSCs recorded in MSNs in which vehicle or MLi-2 (200 nM) were intracellularly perfused through the patch-pipette. (**B)** Box plot and cumulative distributions for mIPSC amplitude in the experimental conditions shown in A (Mann-Whitney test, *p*=0.0006, *U*=65). (**C,D and E)** Box plots showing that MLi-2 does not affect mIPSC decay time (Unpaired Student’s *t-*test, p=0.114, *t*=1.619, *df*=36) frequency (Mann-Whitney test, *p*=0.068, *U*=116.5) and rise time (Unpaired Student’s *t-*test, *p*=0.74, *t*=0.3341, *df*=36). For each parameters vehicle *n/N*=21/6; MLi-2 *n/N*=17/6.

### LRRK2 G2019S impacts on GABAergic transmission

To further investigate the effect of LRRK2 on GABA transmission, we used the technique of “membrane micro-transplantation” in *X. laevis* oocytes (Fig. 2A), which allows the recording of currents from functional receptors^21^ starting from small samples tissue. The GABA-evoked inward current (I_GABA_) in response to GABA (1 mM) was measured in oocytes injected with WT mouse striatal membranes. According to the results obtained from *ex-vivo* mouse brain slices (Fig. 1), the I_GABA_ recorded in oocytes injected with WT striatal membranes was significantly reduced after the inhibition of the kinase activity with LRRK2 inhibitor MLi-2 (I_GABA_ WT: -64.6 ± 8.2 nA *vs* -42.7 ± 5.6 nA before and after MLi-2 incubation, respectively; paired Student’s t-test, *p*<0.0001; not shown). We next assessed the effect of the LRRK2 G2019S mutation on striatal membranes from Lrrk2 G2019S KI mice. I_GABA_ recorded in oocytes injected with LRRK2 G2019S striatal membranes was significantly reduced compared to injected with WT membranes (I_GABA_ WT: -67.1 ± 4.2 nA *vs* I_GABA_ G2019S: -45.7 ± 3.4 nA; Mann-Whitney test, *p*<0.0001; Fig. 2B). Moreover, we analyzed excitatory transmission in oocytes injected with LRRK2 G2019S striatal membranes. The AMPA-evoked current (I_AMPA_) after substrate perfusion (AMPA 20 µM + CTZ 10 µM) was significantly increased in oocytes injected with LRRK2 G2019S striatal membranes (I_AMPA_ WT: -18.3 ± 1.4 nA *vs* I_AMPA_ GS: - 23.07± 1.4 nA; Mann-Whitney test, *p*=0.01; Fig. 2C). In addition, the oocytes injected with G2019S membranes showed a significant increase in NMDA receptor-mediated currents (Glu 200 µM + Ser 10 µM + Gly 10 µM; I_Glu_ WT: -6.7 ± 0.7 nA *vs* I_Glu_ G2019S: - 15.6 ± 1.3 nA; Student’s t-test, *p*<0.0001; Fig. 2D). We further measured the AMPA/GABA ratio and found a significant increase in the LRRK2 G2019S samples (WT: 33.9 ± 1.8 % *vs* G2019S:: 50.2 ± 2.6 %; Student’s t-test, *p*<0.0001; Fig. 2E).

**Figure 2.**
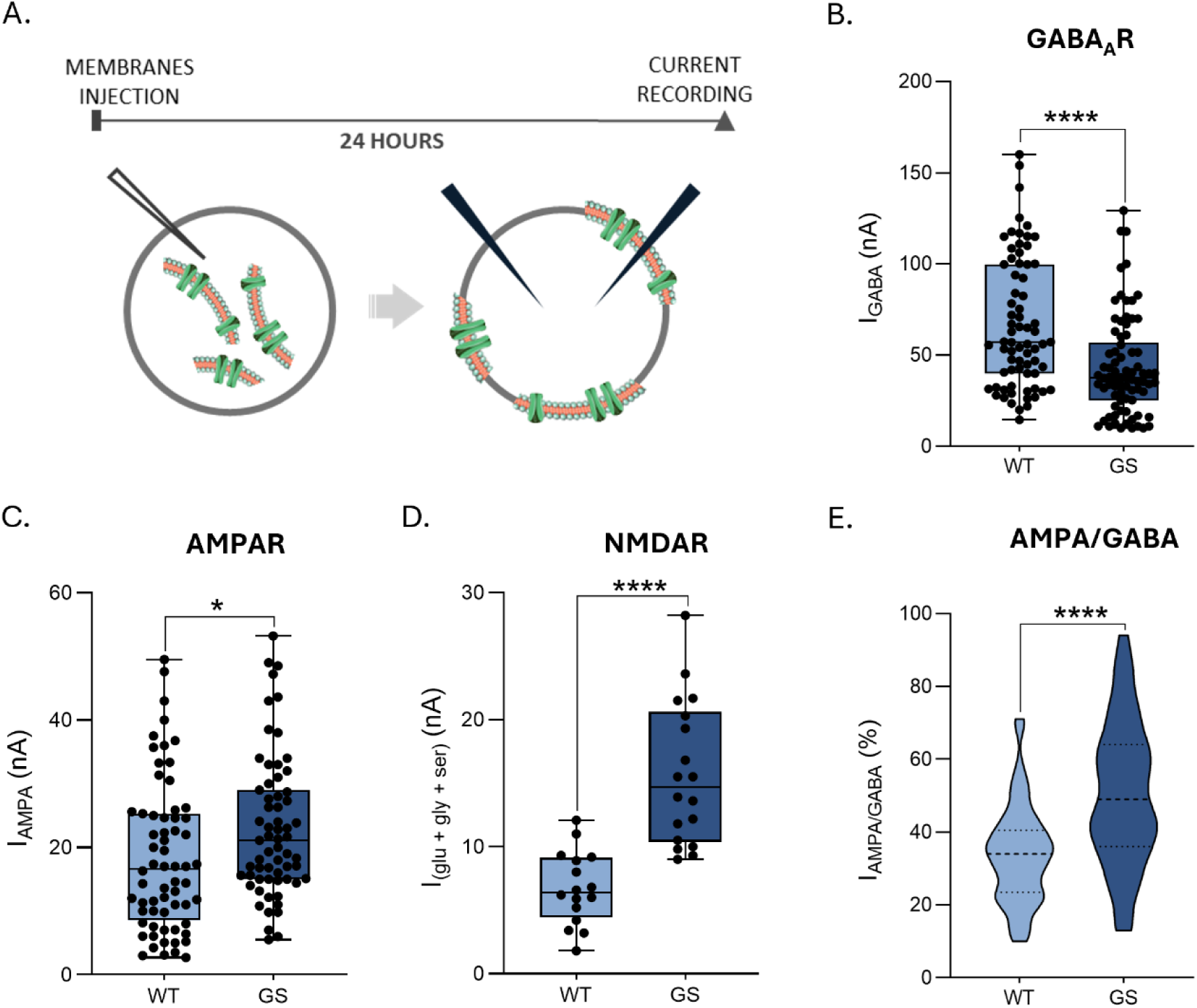
LRRK2 G2019S disrupts the E/I balance. (**A)** Schematic outline of micro-transplantation technique in *X. laevis* oocytes. (**B)** The box plots represent the amplitudes of the GABA current evoked in WT (*n/N*= 71/6) and LRRK2 G2019S (GS; *n/N*=80/6; Mann-Whitney test, *p*<0.0001, *U*=1662) striatal membranes. **(C, D)** AMPA currents evoked in WT (*n/N*=64/4) and LRRK2 G2019S (*n/N*=63/4; Mann-Whitney test, *p*=0.0120, *U*=1497) striatal membranes and glutamatergic current NMDAR-mediated evoked in WT (*n/N*=16/3) and LRRK2 G2019S (*n/N*= 19/3; Unpaired Student’s *t-*test, *p*<0.0001, *t*=5.726, *df*=32) striatal membranes as in B. Ratio AMPA/GABA current measured in WT (*n/N*=53/4) and mutated (*n/N*=55/4; Unpaired Student’s *t-*test, *p*<0.0001, *t*=5.036, *df*=106) membranes expressed in oocytes.

### Functional assessment of GABA_A_ receptors in LRRK2 G2019S striatum

Due to the reduced GABAergic currents observed in LRRK2 G2019S striatal membranes, we characterized the electrophysiological properties of the GABA_A_ receptor. The GABA reversal potential (E_GABA_) did not differ significantly between oocytes injected with membranes from WT and LRRK2 G2019S tissues (E_GABA_ WT: -23.9 ± 0.7 mV *vs* E_GABA_ G2019S: -22.3 ± 1.1 mV; Student’s t-test, *p*>0.05; Supplementary Fig. 1A-B). Furthermore, GABA-evoked responses were dose-dependent in both groups, with comparable EC_50_ values (EC_50_ WT: 103.8 ± 2.5 µM *vs* EC_50_ G2019S: 107.7 ± 2.9 µM; Student’s t-test, *p*>0.05; Supplementary Fig. 1C-D). These findings indicate that the observed reduction in GABAergic currents is not due to altered chloride homeostasis or changes in GABA affinity.

Prolonged GABA application (1 mM, 60 seconds) was performed to measure the current decay value of the GABA_A_ receptor in both conditions (Fig. 3A). The decay time, calculated at 10% of the total current (t_0.1_) of GABA_A_ receptors, was significantly slower in oocytes injected with LRRK2 G2019S samples compared to those with WT membranes (t_0.1_ WT: 4.7 ± 0.2 s *vs* t_0.1_ G2019S: 6.9 ± 0.6 s; Mann-Whitney test, *p*<0.0001; Fig. 3A), suggesting altered GABA_A_ receptor subunit composition in the pathological tissue.

**Figure 3.**
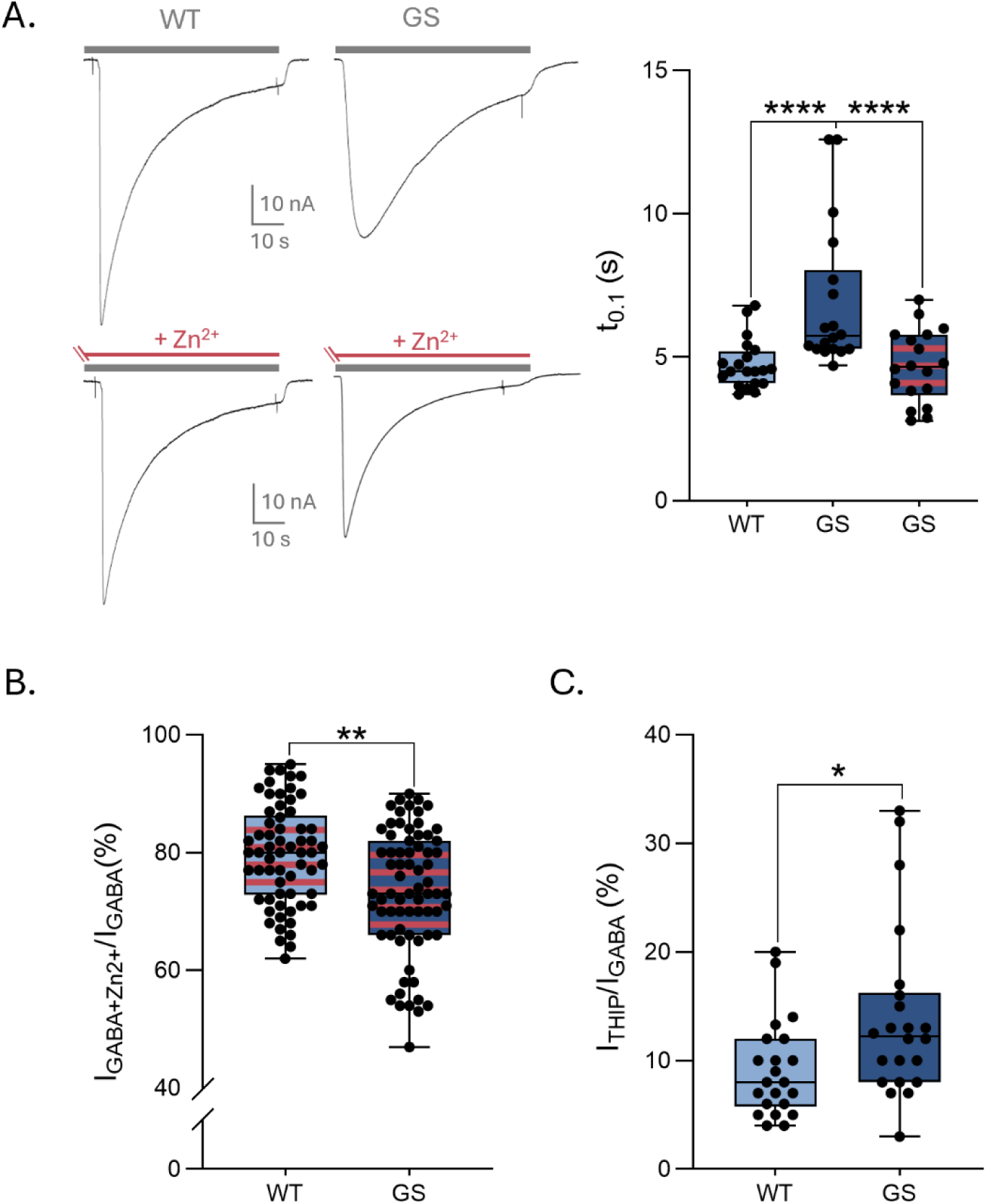
LRRK2 G2019S influences synaptic and extrasynaptic currents. (**A)** *(left)* Representative traces of long pulse of GABA in oocytes injected with WT or LRRK2 G2019S KI mice (GS) striatal membranes before and after (bottom) Zn^2+^ treatment. *(righ)* The box plot shows the decay time measured in oocytes injected with WT tissue (*n/N*=20/3) or with mutated striatal membranes before and after Zn^2+^ treatment (stripped columns) (*n/N*= 18/3, Wilcoxon test, *p*<0.0001). Statistical analysis was performed with ANOVA one-way (ANOVA Kruskal-Wallis test; *p*=0.0003). (**B)** The box plot shows the amplitudes of normalized residual GABA evoked current (%) after Zn^2+^ treatment (stripped columns) in oocytes expressing striatal membrane of WT (*n/N*= 62/6) or LRRK2 G2019S KI mice (*n/N*=71/6; Mann-Whitney test, *p*=0.0019, *U*=1518). Row data were normalized to the total GABA current recorded in each cell. (**C)** The box plots show the amplitudes of THIP evoked current in WT (*n/N*=22/3) or LRRK2 G2019S (*n/N*=22/3; Mann-Whitney test, *p*=0.0121, *U*=136.5) striatal membranes.

To distinguish between synaptic and extrasynaptic GABA_A_ receptors, we treated the oocytes with zinc (Zn²⁺, 40 µM for 120 seconds), which selectively inhibits GABA_A_ receptors lacking the γ_2_ subunit.^22^ Zn^2+^ application reduced GABA-evoked currents in both experimental groups. However, the inhibition was significantly more pronounced in oocytes injected with LRRK2 G2019S membranes, suggesting an impairment of γ_2_-subunit containing GABA_A_ receptors in the pathological tissue (I_GABA_ WT: 79.6 ± 1.08 % *vs* I_GABA_ G2019S: 73.4 ± 1.2 %; Mann-Whitney test, *p*=0.001; Fig. 3B).

Of note, when prolonged GABA application was repeated in the presence of Zn^2+^, the decay time became similar in both conditions (t_0.1_ WT + Zn^2+^: 4.8 ± 0.3 s *vs* t_0.1_ G2019S + Zn^2+^: 4.7 ± 0.3 s; Mann-Whitney test, *p=*0.83; Fig. 2A), indicating that the slower decay time observed in the G2019S condition can be attributed to γ_2_-subunit lacking GABA_A_ receptors. Thus, a parallel experiment was performed to examine the activity of extrasynaptic GABA_A_ receptors. To this end, the oocytes were treated with 4,5,6,7-tetrahydroisoxazolo[5,4-c] pyridin-3-ol (THIP, 50 µM), an agonist of δ-subunit containing GABA_A_ receptors.^23^ The obtained result showed an increase of THIP evoked current in oocytes injected with LRRK2 G2019S striatal membranes compared to the WT counterpart (I_THIP_ WT: -10.9 ± 1.05 nA *vs* I_THIP_ G2019S: -15.4 ± 1.8 nA; Mann-Whitney test, *p*=0.01), corresponding to 9.15 ± 0.94 % and 14.07 ± 1.70 % of the total GABA-evoked current (Fig. 3C) in WT and G2019S conditions, respectively.

### Synaptic GABA_A_ receptor expression in LRRK2 G2019S striatum

Our functional data suggest a reduction of synaptic GABA_A_ receptors in striatal membranes from LRRK2 G2019S mice. To further investigate this issue, we performed western blot (WB) analysis of the γ_2_-subunit of GABA_A_ receptor and gephyrin, a key scaffolding protein of synaptic GABA_A_ receptors. Expression levels were assessed in both total striatal lysates and membrane samples from WT and Lrrk2 G2019S KI mice.

In total striatal lysates, no significant difference in the expression of either gephyrin or the γ_2_ subunit were observed between WT and LRRK2 G2019S samples. In contrast, in LRRK2 G2019S striatal membranes, both gephyrin and the γ_2_-subunit showed a trend towards a reduced expression (Fig. 4B). Although these results did not reach statistical significance (Unpaired Student’s t-test: GPHN: WT vs GS, *p*=0.059; γ_2_-subunit: WT vs GS, *p*=0.104), the selective reduction in membrane fractions, but not in total lysates, suggests a possible involvement of LRRK2 G2019S mutation in the expression or stabilization of GABA_A_ receptors at the synaptic plasma membrane.

**Figure 4.**
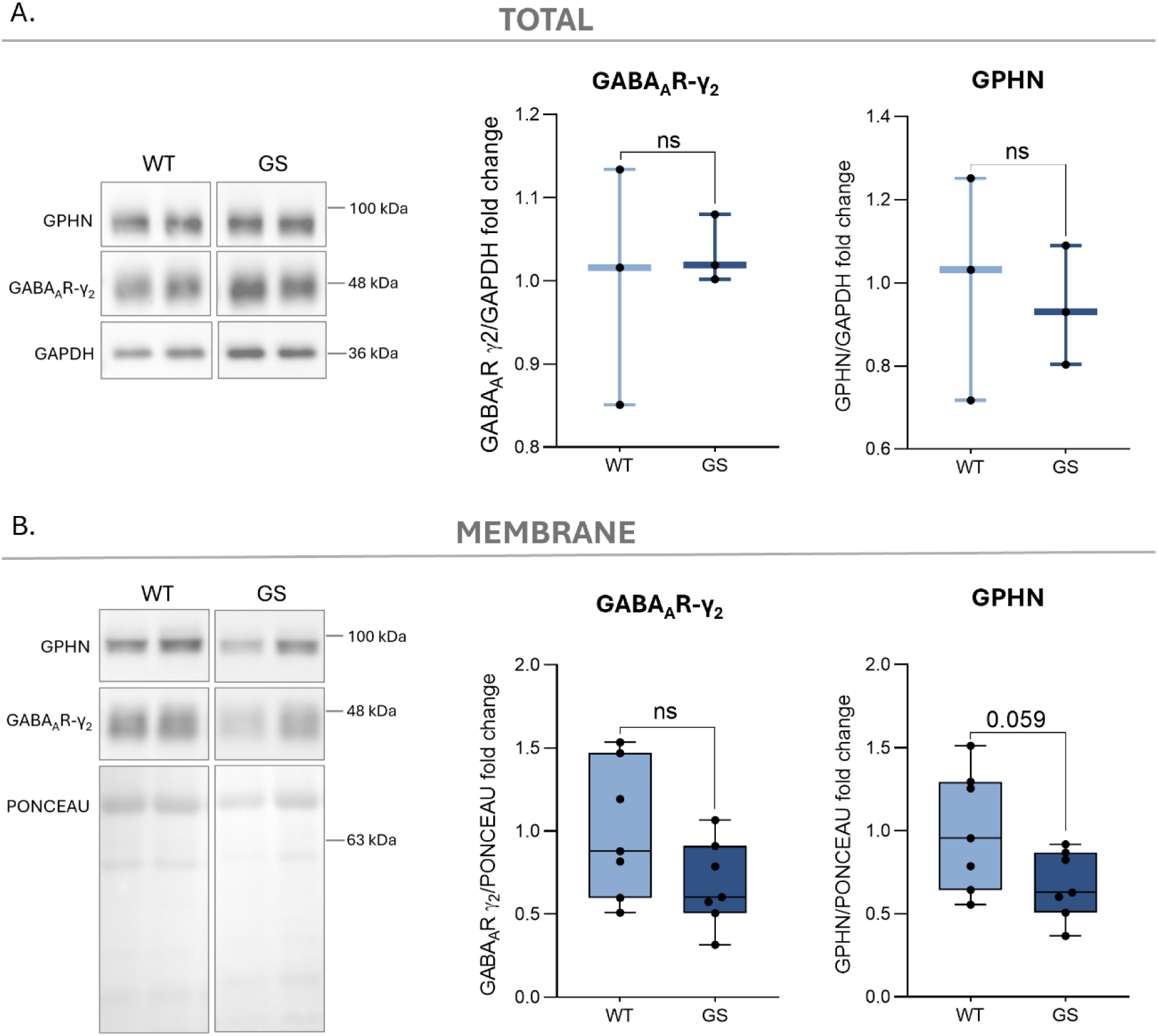
LRRK2 G2019S perturbs membrane levels of synaptic GABA_A_ receptors. **(A)** Representative WB of total striatal lysates of WT and G2019S LRRK2 KI mice. Antibodies against GABA_A_R γ_2_ subunit and gephyrin (GPHN) were used to detect the respective proteins, GAPDH was used as loading control. Box plots show the quantification of normalized γ_2_-subunit (Unpaired Student’s *t-*test*, p*=0.716, *t*=0.390, *df*=4) and GPHN levels (Unpaired Student’s *t-*test*, p*=0.754, *t*=0.335, *df*=4; *n*=3 mice for each group). (**B)** Representative WB analysis of striatal membranes from WT and LRRK2 G2019S KI mice. The ponceau stained membrane was used for normalization. Box plots show the quantification of normalized GABA_A_ receptor γ_2_ subunit (Unpaired Student’s *t-*test*, p*=0.104, *t*=1.758, *df*=12) and GPHN (Unpaired Student’s *t-*test*, p*=0.059, *t*=2.077, *df*=12; *n*=7 mice for each group) levels.

### LRRK2 G2019S functionally interacts with synaptic GABA_A_ receptors

To better understand the functional relation between LRRK2 and the GABA_A_ receptor, we established a system that closely mimics the pathological striatal tissue. To this aim, *X. laevis* oocytes were co-injected with WT striatal membranes and cRNAs coding for either LRRK2 WT or the G2019S mutant (Fig. 5A; See Material and Method Section). This approach allowed to investigate whether overexpression of LRRK2 G2019S could modulate GABA_A_ receptors within the WT striatal membranes. Interestingly, the expression of LRRK2 G2019S caused a significant reduction in GABA-evoked current compared to oocytes expressing the LRRK WT (I_GABA_ WT Striatum + LRRK2 WT cRNA: -66.1 ± 7.2 nA *vs* I_GABA_ WT Striatum + LRRK2 G2019S cRNA: -45 ± 5.1 nA; Student’s *t*-test, *p*=0.03; Fig. 5B), suggesting that the G2019S mutation produces the aberrant phenotype through a tight modulation of the GABA_A_ receptor.

**Figure 5.**
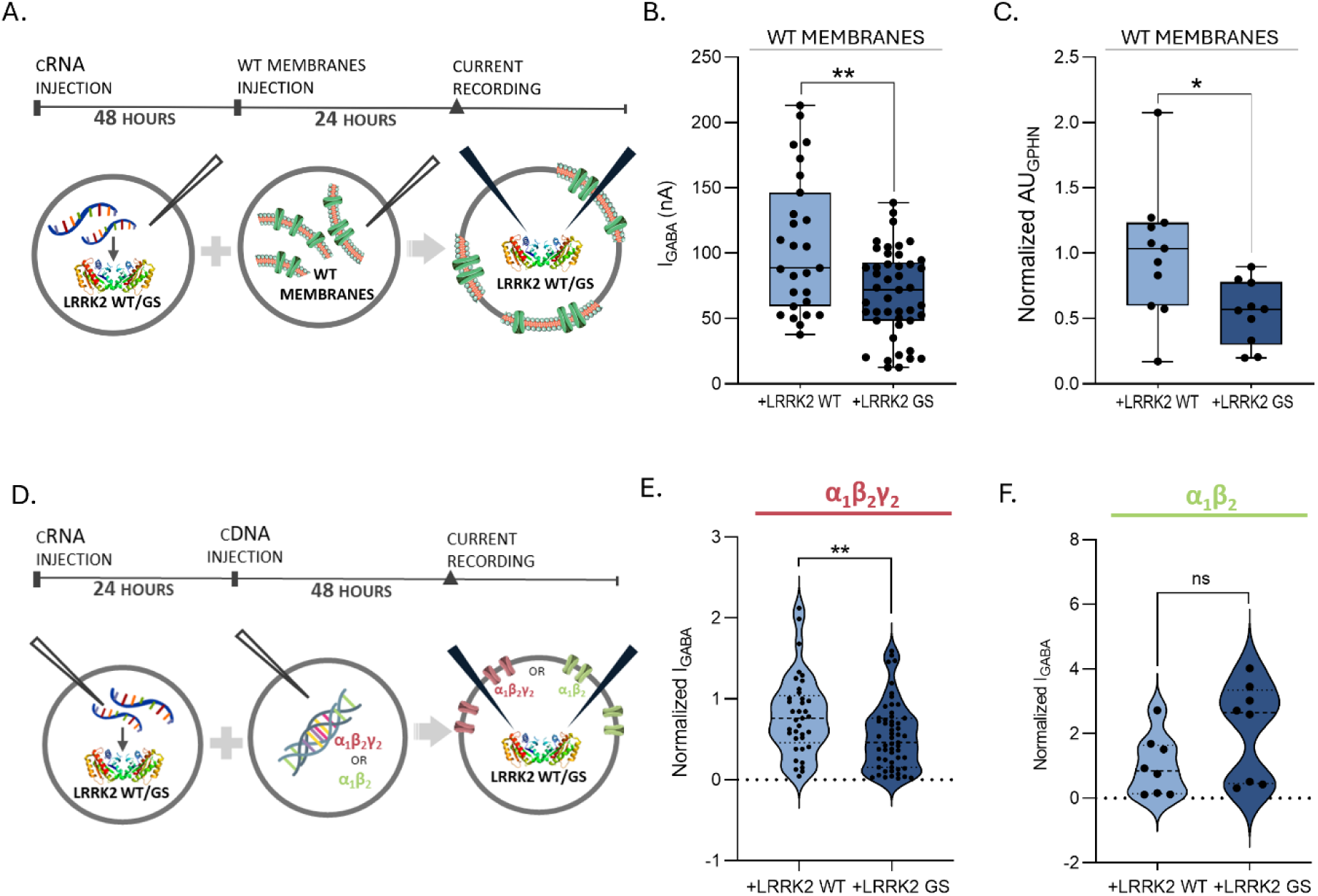
LRRK2 G2019S tightly acts on γ_2_ subunit-containing GABA_A_ receptors. (**A)** Schematic outline of the experimental setup. Oocytes were co-injected with WT striatal membranes plus LRRK2 WT or G2019S cRNA. (**B)** The box plot represents the amplitudes of the GABA evoked current in oocytes co-injected with WT membranes + LRRK2 WT cRNA (*n/N*=27/2) and + LRRK2 G2019S cRNA (*n/N*=45/2; Mann-Whitney test, *p*=0.0094, *U*=386). (**C)** The box plot shows the normalized GPHN amount detected though single oocytes chemiluminescence in oocytes co-injected with WT striatum + LRRK2 WT cRNA (*n/N*=11/2) and + LRRK2 G2019S cRNA (*n/N*=10/2; Unpaired Student’s *t-*test, *p*=0.0152, *t*=2.668, *df*=19). **(D)** Schematic outline of α_1_β_2_γ_2_ GABA_A_ receptor subunits cDNA co-injection with LRRK2 WT or G2019S cRNA in oocytes. (**E)** The violin plot shows the normalized GABA current amplitude recorded in oocytes co-injected with α_1_β_2_γ_2_ GABA_A_ receptor + LRRK2 WT (*n/N*=37/3) or + LRRK2 G2019S (*n/N*=55/3; Mann-Whitney test, *p*=0.0059, *U*=674). The data collected were normalized using the wild-type condition as reference group. (**F)** Violin plot showing normalized GABA-evoked current α_1_β_2_-mediated in co-expression with LRRK2 WT (n/N=8/2) or LRRK2 G2019S cRNAs (n/N=8/2; Unpaired t-test; p=0.08, t=1.82, df=14).

Additionally, the same oocytes were used to measure the amount of gephyrin using the SOC technique upon fixation and permeabilization. Gephyrin levels were significantly reduced in oocytes containing WT striatal membranes co-expressing LRRK2 G2019S, indicating that the mutated kinase disrupts the scaffolding of the synaptic γ_2_ subunit-containing GABA_A_ receptors (Unpaired student’s *t*-test, GPHN WT *vs* GS *p*=0.015; Fig. 5C).

These findings were further supported by a complementary experimental approach aimed at elucidating the functional interaction between LRRK2 G2019S and the canonical synaptic GABA_A_ receptor isoform, α_1_β_2_γ_2_. To this end, oocytes were co-injected with cDNAs encoding the α_1_β_2_γ_2_ GABA_A_ receptor subunits together with the cRNA for LRRK2 G2019S (Fig. 5D). This strategy enabled the overexpression of both proteins, allowing for a more detailed evaluation of their functional interplay.

Co-expression of α_1_β_2_γ_2_ GABA_A_ receptor and LRRK2 G2019S caused a significant reduction in GABA-evoked currents (Mann–Whitney test, *p*=0.0059; Fig. 5E). This reduction was reversed upon treatment with the LRRK2 inhibitor MLi-2 (Wilcoxon test, *p*=0.0039; not shown). To further validate the specificity of the functional interaction between LRRK2 G2019S and the γ_2_-subunit, we used the same experimental system, this time co-injecting cRNA for LRRK2 G2019S together with a binary GABA_A_R receptor isoform lacking the γ_2_-subunit (α_1_β_2_). Under this condition, the presence of either WT or mutant LRRK2 did not significantly alter the amplitude of GABA-evoked currents (Unpaired student’s t-test, p=0.08; Fig. 5F) indicating that the modulatory effect of LRRK2 G2019S requires the presence of the γ_2_-subunit.

### LRRK2, GPHN and GABRG2 are part of a common PPI network

A protein network based on physical interactions was built to identify possible common interactors among GABRG2, gephyrin and LRRK2. The rationale was that common interactors may explain mechanisms by which LRRK2 influences gephyrin and GABRG2 dynamics at the plasma membrane. As shown in Fig. 6A, most of the previous knowledge is related to LRRK2 interactors (and substrates). However, few common interactors are known among these three proteins, as summed up in Table 2 and zoomed in Fig. 6B. This network presents 58 edges and 17 proteins that, beyond being common interactors of GABRG2, LRRK and GPHN, have interconnections among them. The PPI-enrichment p-value is highly significant (*p*<2.2 e^-16^).

**Figure 6.**
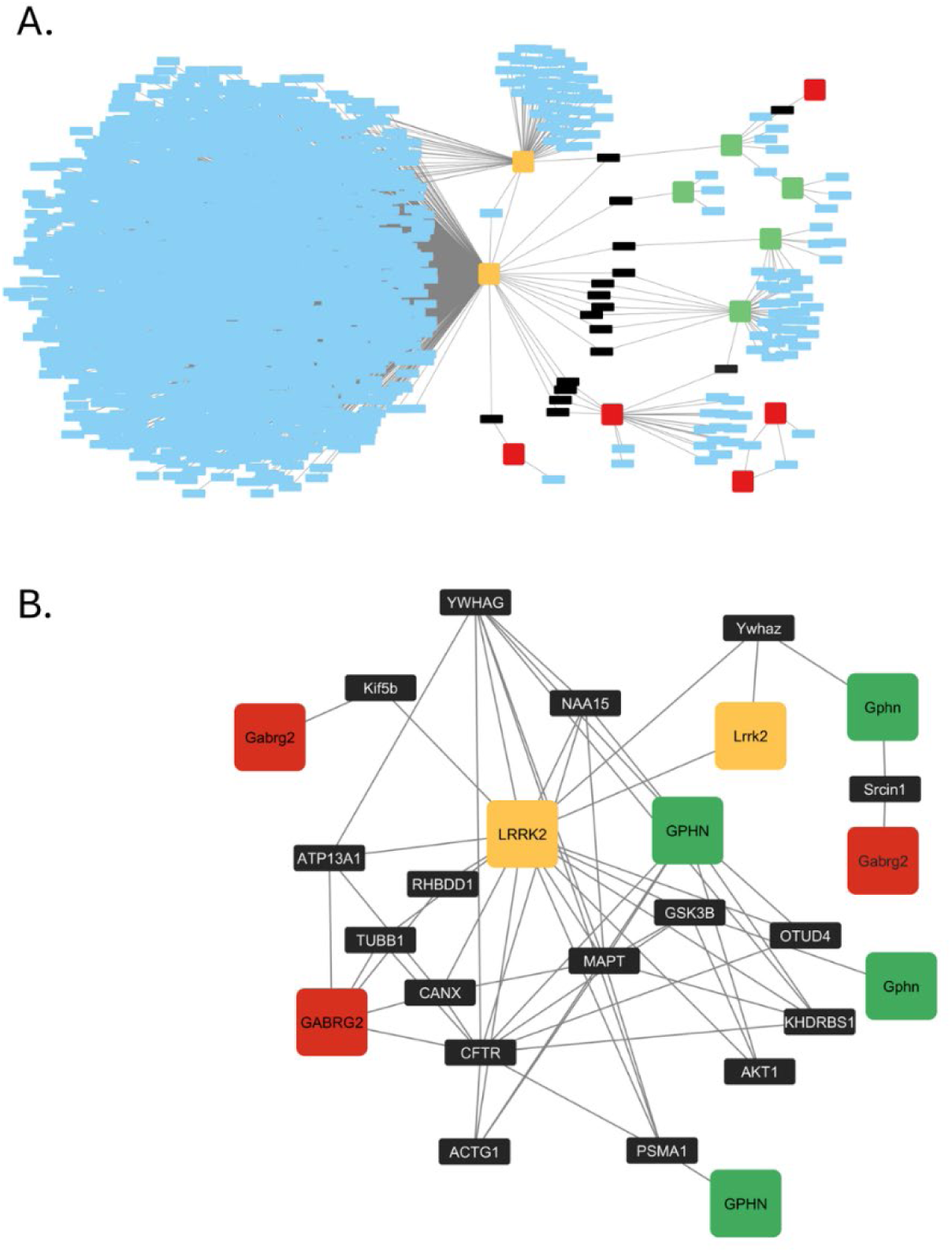
Protein-protein interaction (PPI) network representing all known interactors among GABRG2, Gephyrin and LRRK2. (**A)** All first interactors of GABRG2, Gephyrin and LRRK2. (**B)** Sub-network extracted from A. to focus on common interactors between GABRG2, Gephyrin and LRRK2 and their reciprocal interactions (26 nodes, 58 edges). Red nodes: GABRG2, human; Gabrg2, mouse/rat. Orange nodes: LRRK2, human; Lrrk2, mouse/rat. Green nodes: GPHN, human; Gphn, mouse/rat. Black nodes: common interactors.

**Table 2.** Common interactors among γ2 subunit of GABAA receptor (GABRG2), Gephyrin (GPHN) and leucine rich repeat kinase 2 (LRRK2).

|  |  |  |  |
| --- | --- | --- | --- |
| LRRK2 | and | Gephyrin | YWHAZ, YWHAG, GSK3B, PSMA1, AKT1, KHDRBS1, NAA15, OTUD4, ACTG1, MAPT |
| LRRK2 | and | GABRG2 | ATP13A1, Kif5b, TUBB1, RHBDD1, CANX |
| GABRG2 | and | Gephyrin | Srcin1, CFTR |

## Discussion

The present study reveals an entirely new dimension of LRRK2 biology by uncovering its previously unrecognized role in modulating GABAergic synaptic transmission and revealing a novel mechanism of early circuit dysfunction in Parkinson’s disease.

While LRRK2 has been widely studied in the context of excitatory and dopaminergic signaling, its role in inhibitory circuits remained unexplored. Here, we identify for the first time GABA_A_ receptor a molecular target of LRRK2, establishing a functional link between its kinase activity and postsynaptic GABA_A_R-mediated inhibitory signaling.

We first assess a reduction of mIPSCs recorded in striatal MSNs after postsynaptic inhibition of LRRK2 via MLi-2 that indicates a modulatory role of the kinase on the inhibitory transmission. Expanding our view to the entire synapse (rather than just the postsynaptic side), several factors could contribute to an alteration in mIPSC amplitude. These include a decrease in the quantity of GABA stored within synaptic vesicles,^24^ a reduction in the number of receptors on the postsynaptic membrane^25,26^ and/or modifications to the receptors’ subunit composition or phosphorylation status, which could affect their single-channel characteristics.^27^ Recent studies have shown a reduced presynaptic GABA_B_-mediated inhibition in the motor cortex in PD. Remarkably, this abnormality is not corrected by clinically effective dopaminergic medications, suggesting that it is a non-dopaminergic consequence.^28,29^ However, since LRRK2 inhibition was achieved selectively at the postsynaptic level, our functional data suggest that the reduction of mIPSC amplitude is produced by a decrease in the number/functionality of GABA_A_ receptors at the postsynaptic level. It remains to be addressed whether LRRK2 also affects presynaptic mechanisms of GABA transmission.

Notably, the ex-vivo result was confirmed studying GABA evoked current in X. laevis oocytes transplanted with striatal membranes from wild-type mice before and after MLi-2 treatment. X. laevis oocytes are suitable for studying neurotransmitter receptors and channels via “membrane micro-transplantation,” using minimal amounts of brain tissues. The technique allows for the collection of a large number of cells in which membrane proteins preserve native properties along with their associated accessory proteins.^13,21,30^ As a result, it minimizes the number of animals required to achieve statistically significant data, aligning with the 3Rs principles.

Thus, striatal membranes from LRRK2 G2019S KI mice were microtransplanted in X. laevis oocytes and functionally compared with wild-type counterpart to study the modulation of mutated LRRK2 on GABA signaling.

The results reported here indicate a reduction of GABA current in striatal membranes from LRRK2 G2019S KI mice. Parallel, in the same tissues, we found an increased excitatory current amplitude AMPA and NMDA receptors-mediated, resulting in an altered AMPA/GABA ratio. These findings are consistent with published data showing that cultured cortical neurons from G2019S and R1441C LRRK2 KI rats exhibit larger inward current responses to AMPA and NMDA, along with increased densities of VGLUT1 and PSD-95 puncta.^31^

The functional interaction between LRRK2 G2019S and GABA_A_ receptors has been demonstrated using X. laevis oocytes as a rebuilt system to generate the pathological phenotype starting from WT striatal tissues. The presence of LRRK2 G2019S was able to decrease the GABA current in WT membranes, mimicking a pathological condition. Interestingly, the GABA current reduction is about 30%, similarly to that observed between WT and pathological membranes. Thus, we proved that LRRK2 G2019S tightly modulates GABA_A_ receptors, ruling out the possibility that the reduction in GABA current is merely attributable to chronic events in the tissue.

Notably, both acute pharmacological inhibition of LRRK2 using MLi-2 and its pathological hyperactivation via the G2019S mutation lead to a similar reduction in GABAergic transmission. These convergence effects highlight a critical homeostatic role of LRRK2, where both loss and gain of function disrupt inhibitory signaling. The data indicate a non-linear, dose-sensitive relationship between LRRK2 activity and GABA_A_ receptor function, consistent with a bimodal model in which deviations from physiological kinase levels impair synaptic balance.

However, we found that this mutation does not affect the functionality of GABA_A_ receptors, as indicated by the unchanged GABA reversal potential and GABA affinity parameters. A higher sensitivity to THIP, a δ-subunit-preferring agonist, coupled with a prolonged decay time, suggest an enhanced tonic inhibition in LRRK2 G2019S striatal membranes. At this point, it remains unclear whether in LRRK2 G2019S-associated PD, the increase of tonic inhibition represents a compensatory mechanism to preserve neuronal cell against hyperexcitability.

Conversely, LRRK2 G2019S significantly reduces the synaptic GABA current mediated by γ_2_ subunit-containing GABA_A_ receptors, accounting for the decrease in total GABA current observed in the pathological tissue. Phosphorylation processes and scaffolding proteins precisely control the expression of GABA_A_ receptors on the cell surface. The scaffold protein gephyrin strongly interacts with γ_2_ subunit representing the key protein for the stabilization of synaptic GABA_A_ receptors.^32^ Basically, gephyrin regulates the number of GABA_A_ receptors at synaptic sites through several mechanisms, including gephyrin-gephyrin interaction, gephyrin-receptor binding affinities, gephyrin turnover and synaptic transport. Phosphorylation plays a key role in modulating gephyrin oligomerization, stability, and receptor binding, influencing all aspects of gephyrin dynamics at synapses.^33^

Although no alteration in gephyrin levels is observed in total lysates between WT and pathological samples, we quantified via chemiluminescence a significant reduction of the scaffold protein immediately below the plasma membrane in the presence of LRRK2 G2019S.

Combining protein detection and functional data on phasic currents, our findings suggest that LRRK2 G2019S may alter the exposure of γ_2_ subunit-containing GABA_A_ receptors at the plasma membrane, impacting their trafficking and/or stability by disrupting the scaffold architecture. The unresolved question is whether gephyrin deficiency is a cause or consequence of reduced GABA current.

However, our data indicate that synaptic GABA_A_ receptors and specifically γ_2_-subunit are a key molecular factor in LRRK2 G2019S-related PD. The reduction in GABA-evoked currents observed upon co-expression of LRRK2 G2019S with the full α_1_β_2_γ_2_ receptor isoform, combined with the absence of effect when the γ_2_-subunit is missing, demonstrates that the pathological action of LRRK2 is selectively γ_2_-dependent.

It is reasonable to state that LRRK2 G2019S consistently modulates GABA_A_ receptors independently of other pathological hallmarks commonly associated with the progression of PD, such as loss of dopaminergic neurons, DA depletion or altered co-release of GABA and DA. In our experimental system, the G2019S mutation alone is sufficient to induce GABAergic deficits regardless of the synaptic input to the MSNs. These findings point to a cell-autonomous mechanism by which LRRK2 G2019S intrinsically alters inhibitory signaling. Importantly, these effects were observed in dopamine-independent systems, reinforcing the hypothesis that LRRK2 may act upstream of dopaminergic neurodegeneration. This observation is consistent with previous data showing early LRRK2 activation prior to nigrostriatal degeneration.^34^ Thus, our data therefore suggest that GABAergic dysfunction could represent an initiating event in PD pathogenesis, rather than a downstream consequence of dopamine loss.

Nevertheless, we suggest that additional proteins may be involved in the functional link between the kinase and GABA_A_ receptor, contributing to the complexity of the underlying mechanism. The PPI network generated querying γ_2_ subunit of GABA_A_ receptor, LRRK2 and Gephyrin evidenced several common interactors among these three proteins, pointing out possible enzymes and scaffold proteins that may play a role in the effect of LRRK2 G2019S on GABA_A_ receptor. Moreover, the PPI network underlined that these three proteins collaborate for the organization of the synapse, being the PPI enrichment p-value highly significant. The phosphorylation role in regulating the entire process may be crucial. LRRK2 interacts with 14-3-3 family proteins, involved in regulating cellular signaling and proteins trafficking, in a phosphorylation-dependent manner. This interaction modulates the stability of LRRK2 and its kinase activity, influencing the membrane delivery and proteins recycling. A disruption of binding between LRRK2 and 14-3-3 protein has been found when pathogenic mutation on LRRK2 gene, such as the substitution G2019S, occurs.^35^ Interestingly, Wen and colleagues^36^ demonstrated that 14-3-3 acts as a chaperon protein that forms a cargo adaptor complex, together with huntingtin-associated protein 1 (HAP1), regulating the GABA_A_ receptor expression at plasma membrane and, consequently, the inhibitory post-synaptic current.

Furthermore, it has been reported that GSK3β influences the phosphorylation state of gephyrin and, consequently, the membrane exposure of GABA_A_ receptor and related inhibitory current.^37^ This pathway may be mediated by LRRK2, which directly interacts with GSK3β and enhances its kinase activity.^38^ Moreover, LRRK2 also phosphorylates Akt1,^39^ which subsequently interacts with gephyrin,^40^ further linking these molecular signaling events.

It is known that Ca^2+^ entry via NMDA receptor leads to synaptic GABA_A_ receptors displacement mediated by activation of calcineurin, which dephosphorylates Serine 327 of γ_2_-subunit resulting in reduced synaptic residency of GABA_A_ receptors.^41^ This process is aligned with the hyperactivation of NMDA receptors observed in LRRK2 G2019S striatal tissue, and is further reinforced by evidence that LRRK2 physically interacts with RCAN1,^42^ a key regulator of calcineurin. In addition, LRRK2 itself has been shown to interact with calcineurin,^43^ further supporting the involvement of this signaling pathway.

The combination of chronic events in native tissue and the tight modulation of LRRK2 on GABA_A_ receptors, here reported, is the final scenario which we have investigated in the present work.

This study brings out the novel role of LRRK2 in modulating inhibitory signalling by functionally interacting with GABA_A_ receptors localized at synaptic area. The hyper-excitability and the low GABAergic tone responsible for neuronal network damages at striatal synapses are two pathological conditions in which LRRK2 G2019S actively takes part.

In a broader context, our data indicate a pioneering pathogenic mechanism by which the striatal neurons might be affected at the early stage of the neurodegenerative process. Thus, by specifically targeting the cell-autonomous effects of LRRK2 G2019S, therapeutic interventions could rebalance the E/I ratio aimed to contrast or delay the progressive neurodegeneration in PD patients.

Beyond advancing our understanding of early PD pathophysiology, these findings carry significant therapeutic implications. By identifying a previously unrecognized functional axis driving early inhibitory imbalance, mechanistically independent of dopaminergic neuron loss, we open new avenues for interventions targeting the prodromal or preclinical stages of the disease. Strategies aimed at modulating GABA_A_ receptor expression, stabilizing receptor–scaffold interactions, or restoring E/I balance may help delay or mitigate downstream neurodegeneration.

This work adds a critical layer to the emerging picture of early PD, positioning inhibitory dysfunction as a promising target for disease-modifying approaches. Furthermore, given the growing evidence implicating LRRK2 in other brain disorders, uncovering its function in GABAergic regulation points to potential targets for cross-disorder therapeutic strategies beyond PD.

## Data availability statements

The data underlying this article will be shared on reasonable request to the corresponding author.

