## Supplementary Fig. 1 A-B and Fig. 1 C-D for "LRRK2 G2019S disrupts GABAergic signaling and shifts excitatory/inhibitory balance in the striatum"

**Supplementary Figure 1.**

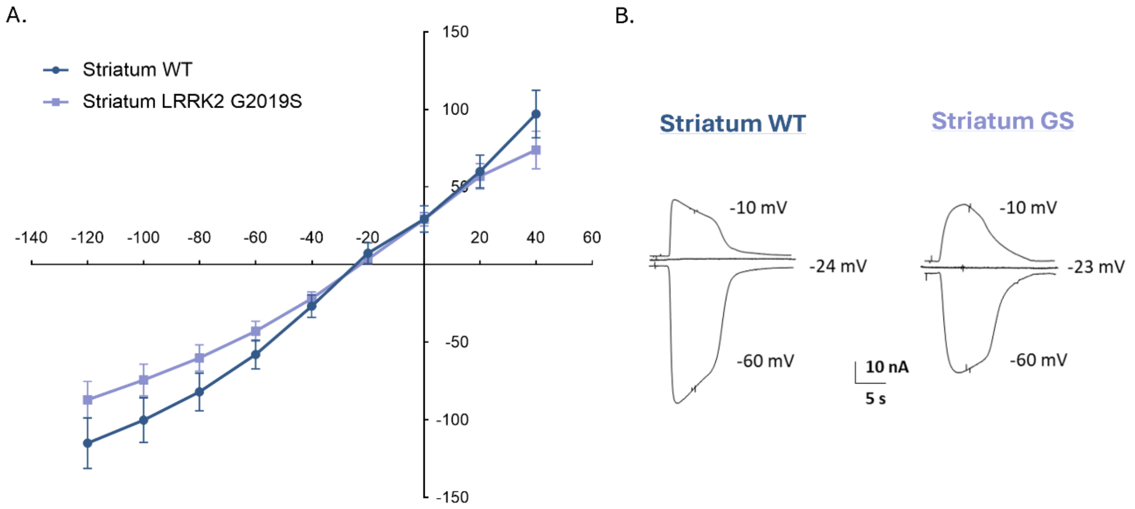


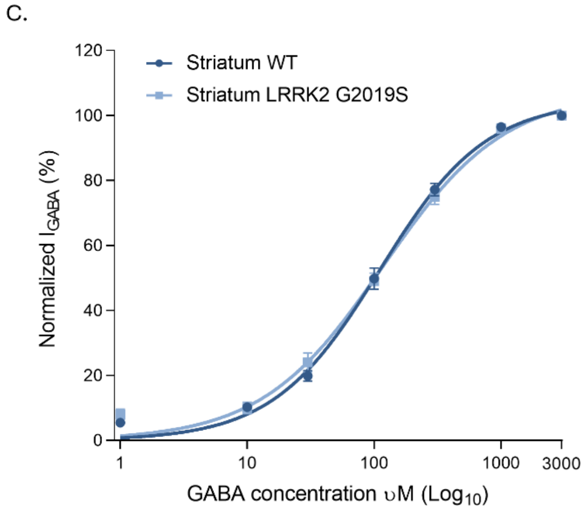


D.

| | $K_{0.5}$ (uM) | Hill coefficient |
|---|---|---|
| **Striatum WT** | 103.8 ± 2.5 | 1.04 |
| **Striatum LRRK2 G2019S** | 107.7 ± 2.9 | 0.96 |

**Supplementary Figure 1. LRRK2 G20129S does not affect $GABA_A$ receptor functionality. (A)** I-V relationships in oocytes injected with striatal membranes of LRRK2 WT (*n/N*= 21/4) or G2019S KI mice (*n/N*= 19/4). The dots represent the mean current amplitude ± SEM at corresponding holding potentials from -120 mV to +40 mV. **(B)** Representative traces of inward, null and outward GABA response in oocytes injected with WT or GS striatum. **(C)** GABA dose-current response relationships. Points represent mean normalized $I_{GABA}$ ± SEM from oocytes injected with WT and LRRK2 G2019S striatal membranes (*n/N*=14/3). **(D)** The dose-response curves were fitted using the Hill equation. The obtained $GABA_AR$ kinetic parameters are reported in the table.
